# TRPA1: A Potential Prognostic Indicator for Oral Squamous Cell Carcinoma

**DOI:** 10.1101/2021.11.29.470496

**Authors:** Rajdeep Chakraborty, Amara Jabeen, Honghua Hu, Charbel Darido, Karen Vickery, Shoba Ranganathan

## Abstract

**Introduction:** Transient receptors are related to oral cancer pain. Previously capsaicin (TRPV1 agonist) was shown to induce cell death in oral cancer cells. We hypothesised that these receptors are present in oral cancer.

**Method:** We examined the presence of cannabinoid receptors (CB1 and CB2) and targets (TRPV1, TRPA1, Ca_V_ 3.1, Ca_V_ 3.2, Ca_V_ 3.3) via quantitative polymerase chain reaction (qPCR) in oral cancer cells SCC4, SCC9, SCC25, Cal27, and normal oral cell line OKF6.

**Result:** Cannabinoid receptors are absent in all the cell lines, while TRPA1 is only present in normal cells, but absent in all the oral cancer cell lines. Voltage-gated calcium channels are present in all the cell lines.

**Conclusion and Future Aspects:** TRPA1 could be the possible future prognostic indicator of oral squamous cell carcinoma. Future functionality assays could use precancerous cell lines to follow the loss of TRPA1.

---

Oral cancer is the sixth most common cancer worldwide and ranks in the top three of all cancers in India.^[1]^ Oral cancer pathogenesis has been associated with the aberrant expression of oncogenes such as *ras, c-myc, bcl-1*, and epidermal growth factor receptor (EGFR/*c-erb 1*).^[2]^ Cannabinoids have been shown to inhibit epidermal growth factor-induced proliferation and metastasis of breast cancer cells.^[3]^ Cannabinoids have all been shown to affect cannabinoid receptors such as CB1 and CB2 and cannabinoid targets such as TRPA1 (transient receptor potential ankyrin 1), TRPV1 (transient receptor potential vanilloid 1), Ca_v_ 3.1, 3.2, and 3.3 (voltage gated calcium channel).^[4]^ We have recently shown that capsaicin, an agonist of TRPV1, can induce cell death in oral cancer cells.^[5]^ Also, transient receptor channels are known to be related to pain associated with oral cancer.^[6]^ Therefore, we hypothesised that drugs that can affect the function of these receptors can inhibit the progression of oral cancer. Our first step was to establish the presence of these receptors in oral cancer cells via quantitative polymerase chain reaction (qPCR). Four oral cancer cell lines, SCC4 (ATCC CRL-1623), SCC9 (ATCC CRL-1629), SCC25 (ATCC CRL-1628) and Cal27 (ATCC CRL-2095) and a normal oral cell line OKF6 (CVCL_L222) were used in this study, with bioethics approval from the Macquarie University Biosafety Committee (“Mammalian Cell Culture” #5215). Molecular profiling of SCC4, SCC9, SCC25, Cal27, and OKF6 previously shown that they can be used as preclinical models for oral cancer translational research.^[7]^ Following ATCC guidelines, Cal 27, SCC4, and SCC9 were cultured using Dulbecco’s Modified Eagle Medium (DMEM) (Gibco, Cat #11960-044) plus 10% foetal bovine serum (FBS, Life Technologies, Cat #10099141) plus 1% penicillin (P)/streptomycin (S) (Life Technologies, Cat #15140-148). To culture OKF6 and SCC25, we used keratinocyte serum-free medium (KSFM, Life Technologies Cat #1700504) plus growth factors (Gibco: human recombinant epithelial growth factor (EGF), Cat #10450-013 and bovine pituitary extract (BPE, Cat #13028-014). The medium for all cell lines was changed twice a week. Each 75 cm^2^ flask of cells acted as a biological replicate, with each biological replicate split into three technical replicates. Biological replicates i.e. flasks were throughout from three different cell passages and conducted on different days (or at different time points). To conduct qPCR, we first selected the probes, that was based on human genome maps, from Thermo Fisher Scientific (https://www.thermofisher.com/order/genome-database/). Only probes with suffix_*m* were selected because they span exon–exon junctions, helping to reduce the chances of genomic DNA (gDNA) amplification. Probes selected for this experiment are TRPC subfamily V member 1 (TRPV1) TaqMan Gene Expression Assay (FAM) Assay ID: Hs00218912_m1; TRPC subfamily A, member 1 (TRPA1) TaqManGene Expression Assay (FAM) Assay ID: Hs00175798_m1; Calcium voltage-gated channel subunit α1 G (Ca_v_ 3.1) TaqMan Gene Expression Assay (FAM) Assay ID: Hs00367969_m1; Calcium voltage-gated channel subunit α1 H (Ca_v_3.2); TaqMan Gene Expression Assay (FAM) Assay ID: Hs01103527_m1; Calcium voltage-gated channel subunit α1 I (Ca_v_ 3.3) TaqMan Gene Expression Assay (FAM) Assay ID: Hs01096207_m1; Glyceraldehyde 3-phosphate dehydrogenase (GAPDH) TaqMan Gene Expression Assay (FAM) Assay ID: Hs99999905_m1; Cannabinoid receptor 1 (CB1) TaqMan Gene Expression Assay (FAM) Assay ID: Hs00275634_m1; and Cannabinoid receptor 2 (CB2) TaqMan Gene Expression Assay (FAM) Assay ID: Hs00361490_m1. Following the manufacturer’s instructions, the TaqMan Fast Advanced Cell-to-CT kit was used to conduct qPCR. To conduct qPCR, four positive control cell lines previously reported to express specific receptors and targets were used, being MCF-7 (positive control of TRPV1, Ca_v_ 3.2, and Ca_v_ 3.3); HT-29 (positive control of TRPA1); PBMC (positive control of CB2) and SH-SY5Y (positive control of CB1 and Ca_v_ 3.1).^[8],[9]^ cDNA was synthesised from 50,000 cells per well of a 96-well plate for all the oral cancer, normal and positive control cell lines. QuantSudio RealTime PCR Software v1.3 was used to collect the results from the PCR reactions. Amplification (ΔRn v. cycle; Rn v. cycle; Ct v. well number), multicomponent (FAM, ROX), raw data and gene expression plots were obtained. The delta−delta C_t_ formula (2^-ΔΔCt^) was used to calculate the relative fold gene expression of cannabinoid receptors and targets. C_t_ is the threshold cycle. The formula used was as follows:

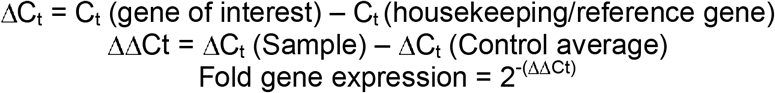

After normalising against the gene expression of the reference gene, *GAPDH*, and calibrating the gene expression in each cell line (five passage numbers at different time points plated in triplicate; i.e. *n* = 15) against that in the normal oral cell line, OKF6, the relative fold gene expression of the cannabinoid receptors and targets for each cell line was studied. Cannabinoid receptor 1 (*CNR1*) and cannabinoid receptor 2 (*CNR2)* expressions are absent in all oral cancer and the normal oral cell line (Table 1). Interestingly, TRPA1 expressions are only present in normal oral cell line (Table 1). None of the oral cancer cell lines studied here showed the presence of TRPA1 expression. Ca_v_ 3.1, Ca_v_ 3.3, and TRPV1 expressions are present in all the oral cancer and normal oral cell line. Incidentally, we have recently shown that capsaicin, a TRPV1 agonist, is effective against oral cancer cell proliferation.^[5]^ Also, Ca_v_ 3.2 expression is only present in SCC25 (Table 1).

**Table 1:**
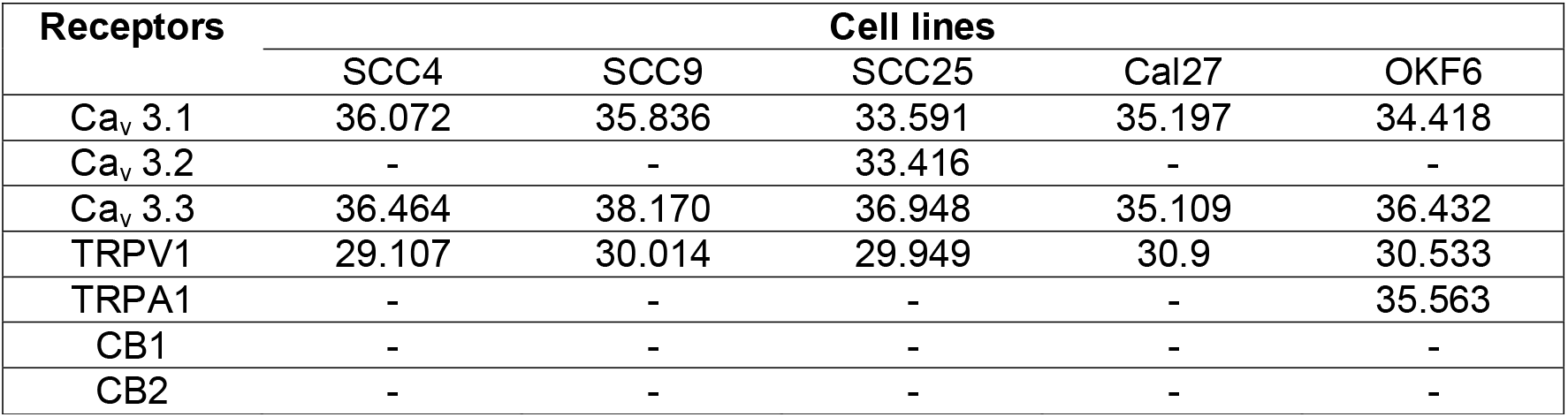
Mean C_t_ values of each receptors.

Ever since the discovery of TRPV1 receptor by the team of Prof David Julius in 1997,^[10]^ that eventually helped him to win The Nobel Prize in Physiology or Medicine 2021, a plethora of experiments worldwide found various other transient receptor channels in human body. They are mainly related to pain perception, while a few are implicated in cancer cell proliferation. Transient receptors affect cellular calcium homeostasis, which plays a pivotal role during the proliferation of cancer cells.^[11]^ Oral cancer is no exception. Interestingly, TRPA1 has been previously shown to be related to oral cancer pain attributed to calcium entry into the cells, facilitated by TRPA1.^[6]^ However, we found that none of the oral cancer cells studied here showed TRPA1 expression, whereas it is present in the normal oral cell. This result gives us a strong indication that TRPA1 could become a future prognostic indicator of oral squamous cell carcinoma. It would be interesting to get a clear picture of the gradual loss of TRPA1 from normal to cancerous state of the cell. Therefore, in future TRPA1 expressions of cells from precancerous lesion (leukoplakia) or at different stages of dysplasia could be investigated.

Cannabis, in marijuana, is not associated with head and neck squamous cell carcinoma or oral squamous cell carcinoma.^[12]^ Our results are concordant with this study, with no oral cancer cell line showing the presence of cannabinoid receptors.

Also, we found, for the first time, the expression of voltage-gated calcium channels (VGCC) in oral cancer cells (Ca_v_ 3.1, 3.2, and 3.3). Serendipitously, we recently found that ML218 HCl (a VGCC drug) is effective in inhibiting bacterial antigen-induced Cal 27 oral cancer cell proliferation.^[13]^ Therefore, future functionality assays of TRPA1 and VGCC via Calcium 5 assay and patch clamp assay, respectively, are essential to develop new therapies for the management of oral cancer via TRPA1/VGCC.^[14],[15]^ Careful excision of the precancerous/cancerous lesion is necessary to avoid neural cell contamination for the determination of TRPA1 expression.

The inherent limitation of the project is the use of oral cancer cell lines from Caucasian subjects. Epigenetically, oral cancer cell lines of Asian or Indian subcontinent origin may differ.^[16]^ Given the preponderance of oral cancer among the Indian subcontinent population,^[1]^ oral cancer cells of Indian subcontinent or Asian origin should also be used in future research.

Finally, we would like to conclude that this report provides initial indications of TRPA1 as a possible future prognostic indicator for oral squamous cell carcinoma. We also report the complete absence of cannabinoid receptors in oral cancer and normal cells. Also, we strongly encourage further research in the field of voltage-gated calcium channels in head and neck cancer.

The project was funded by the Sydney Vital Translational Cancer Research Award Round 9. Rajdeep Chakraborty is the recipient of the International Macquarie University Research Excellence Scholarship (iMQRES).

We would like to thank Dr Marina Junqueira Santiago and Thermo Fisher Scientific for their technical advice during the project.

## Notes

### Competing Interest Statement

The authors have declared no competing interest.

